# Benefits and Challenges of Integrating a Generative AI Assisted Reading Guide in an Undergraduate Journal Club Assignment

**DOI:** 10.64898/2026.02.26.708236

**Authors:** Ashley Ringer McDonald, Anne V. Vázquez

## Abstract

Developing scientific reading skills is critical for undergraduate STEM students due to scientific literature’s unique formatting and use of specialized jargon. Generative AI tools such as ChatGPT offer students the ability to ask questions about what they are reading interactively. Previously, we reported the development of a ChatGPT-assisted reading guide that combined structured, active reading strategies with using ChatGPT to clarify unfamiliar words and concepts in real time. In the initial study, undergraduates found the use of the ChatGPT-assisted reading guide helpful in their understanding of an abstract and introduction of a journal article. Here, the ChatGPT-assisted reading guide was used in a journal club assignment for an undergraduate chemistry course. ChatGPT transcripts were analyzed for common types of interactions, and students were surveyed about their experience. Overall, students reported that using the ChatGPT-assisted reading guide was helpful in understanding the article and helped them have more productive class discussions. However, some students also expressed skepticism about using AI tools, citing concerns about accuracy of AI-generated information and the effect of using AI on their own learning.

## Introduction

Reading scientific literature is a crucial skill for scientists, but how and to what extent it is taught in the undergraduate STEM curriculum is variable. Scientific literature poses unique challenges for learners, as these papers have unique formats, contain unfamiliar jargon, and require the interpretation of data and graphs/figures within the context of the paper.^1^ Further, it has been shown that there is a disconnect between undergraduates’ perceived ability to read literature and actual comprehension,^2,3^ with undergraduate students frequently struggling to identify hypothesis and supporting evidence for conclusion in papers^3^. Poor reading ability has been associated with increased academic anxiety, which can impact persistence in STEM fields.^3^ Likewise, studies have also shown that active engagement with scientific literature increases sense of belonging in STEM,^1,4-7^ and it is therefore imperative to develop pedagogical tools for teaching undergraduate students to actively and effectively read scientific literature.

There have been successful interventions to improve scientific reading skills for undergraduate STEM students. One guided curriculum uses “figure facts” to focus on figures and the experimental data used to create them. This curriculum led to gains in data interpretation skills and confidence.^7^ Another curriculum done in a workshop setting used scaffolded worksheets focused on identifying hypotheses and conclusions drawn from experimental data, leading to improvement in confidence.^3^ A different curriculum involved students actively peer reviewing preprints as a means to improve understanding of scientific literature. The authors of this study noted gains in literacy skills and sense of belonging in STEM.^8^

While the studies described above make significant gains in helping students engage with scientific literature, the issue of unfamiliar language or jargon remains. Encountering unfamiliar terms can be a major barrier for diverse learners with different language backgrounds, and students often simply skip or skim information they do not understand, leading to poor learning outcomes.^2^ Generative AI’s natural language minimizes these barriers for students,^9-11^ allowing them to obtain definitions for unknown jargon in real time, and within the paper’s and learner’s context. A study by Haraldsrud and Odden^12^ investigated productive uses of generative AI in the context of chemistry computational modeling. Using the theoretical framework of distributed cognition, in which cognitive processes are understood to extend beyond the individual to their interaction with others, tools, and the environment, the authors investigated ways students used generative AI in solving chemistry computational problems. Productive themes in interactions with AI were found to be “leveraging AI to retrieve and clarify information” and “using GenAI to test and critique ideas.” Nonproductive themes in interactions with AI were found to be “outsourcing thinking to the AI” and “when GenAI output exceeds student understanding.” In their study, approximately two-thirds of student interactions with AI were coded to be productive, and the authors suggest that to encourage the productive use of AI, students should use AI only after engaging in their own analysis, give AI sufficient context when submitting a query, and that interactions with AI should be dynamic and interactive in nature.^12^

We believe combining the use of generative AI with proven active reading strategies can improve undergraduate STEM students’ ability and comfort in engaging with scientific literature by encouraging the productive types of interactions discussed in the paper by Haraldsrud and Odden.^12^ Here, we report on the use of a ChatGPT-assisted reading guide to support students in a journal club assignment in a chemistry seminar course. The ChatGPT-assisted reading guide combines active reading strategies with the use of ChatGPT to clarify unknown terminology in real time as the student reads the journal article, with the goal of encouraging productive use of generative AI to help students engage with scientific literature.

### The ChatGPT-assisted reading guide

To improve active engagement with scientific literature, in a previous paper we described the development of a ChatGPT-assisted reading guide that combines active reading strategies with the use of ChatGPT to assist students in clarifying unknown terms and concepts in real time.^13^ The active reading and notetaking strategies in the reading guide are adapted from McGuire’s book *Teach Students How to Learn*.^14^ The ChatGPT-assisted reading guide consists of four steps to guide students in using the chatbot to supplement active reading. The full reading guide can be found in Supplemental Materials, and is briefly described here.

1. Previewing the paper: Analyze the title, abstract, headings, and figures to gain meaning
  a. Title: Identify key words that show the paper’s main focus
  b. Abstract: Identify key findings and the general format of the abstract (i.e., significance, problem to be addressed, approach, and insights)
  c. Headings and Figures: Skim text headings and figures to anticipate questions the paper will answer
2. Guided reading of the paper: Highlight key text with an erasable method and refine or edit highlighting as you read; make margin notes of significant points; write extended notes in a separate notebook as needed.
3. Using ChatGPT for clarifications: Use ChatGPT to help clarify unfamiliar terms, concepts, or methodologies.
  a. Introduction: Introduce yourself with academic background, research experience, and any other pertinent information
  b. Paper introduction: Inform ChatGPT about the paper, including the title and general field of the paper
  c. Questions: Ask questions to ChatGPT and ask follow-up and clarifying questions as needed.
4. Post-reading summary: Review notes and the ChatGPT transcript to write a brief summary of the paper to synthesize ideas and serve as a record of the paper.

In an initial study, students engaged in directed undergraduate research were asked to use the ChatGPT-assisted reading guide to read the abstract and introduction of a paper. Pre- and post-surveys were administered to assess students’ experience with scientific reading, as well as their experience using the protocol. Anonymized ChatGPT transcripts were also collected and student interactions with ChatGPT were coded for common themes. Students generally had a positive experience using the ChatGPT-assisted reading guide and felt it helped them to engage with the article more effectively.

### Context of the class

To examine the utility of the reading guide in a larger context, we conducted a follow-up study with undergraduate students enrolled in a second-year seminar course at Cal Poly San Luis Obispo. The course is a one-unit course required for all chemistry and biochemistry majors that meets once a week for 50 minutes; the enrollment in the course in the term this study was conducted was 29 students. The course learning objectives include resume building, developing professional communication skills, and learning to read primary literature in chemistry and biochemistry. Traditionally, the course included a “journal club” component where students read an assigned paper on their own and discussed the paper in small (4-5 people) groups with their peers during class. The paper is selected by the instructor and everyone in the class is assigned to read the same paper.

In our study, the class included a “journal club” reading assignment in two different weeks. In the first iteration, the students read the assigned paper on their own, without any specific reading instructions and or using a large-language model. We did not give students specific instructions about using other resources; in prior iterations of the course, the students said they had looked up unfamiliar terms or concepts using an internet search engine. Since this was not expressly disallowed, it is probable the students did similar things, but we did not ask the students about this specifically in the survey. The students then had a small group discussion with their peers for approximately 25 minutes, followed by an approximately 15 minute full-class discussion led by the instructor. In the second iteration, students were asked to read the paper using the reading guide described above, including the specific instructions about interacting with the LLM.

Students submitted the transcripts from their LLM conversation. Additionally, the students still had an in-class discussion with their journal club groups during the next class meeting. The discussion groups were the same for both weeks of discussion.

All students enrolled in the course completed the traditional and LLM-assisted journal club assignments as required components of the course. Participating in the survey about their experience and having their ChatGPT transcript analyzed as data for the study was optional. The survey link, which included a consent form to participate in the study, was posted on the course learning management page. The instructor then collected the ChatGPT transcripts for the students who agreed to participate in the study and anonymized them before they were coded by the research team. This procedure was approved by the institutional review board at both St. John’s University and Cal Poly San Luis Obispo.

### Coding ChatGPT transcripts

ChatGPT transcripts were collected and anonymized, and 21 complete transcripts were used for analysis. First, the number of interactions that the student initiated with ChatGPT were tallied. An interaction was counted as any time the student typed in the ChatGPT conversation. The mean number of interactions was 4.8 (*SD* 2.49), with a range of 1 to 10 interactions.

The types of interactions were coded using a hybrid coding scheme where an initial set of five themes from the first study were used,^13^ and then iterative open coding was used to identify any new themes in this study. All themes from the initial study were observed, as well as two emergent themes. The codes and the frequency of use is seen in **Figure 1** and are discussed below.

**Figure 1.**
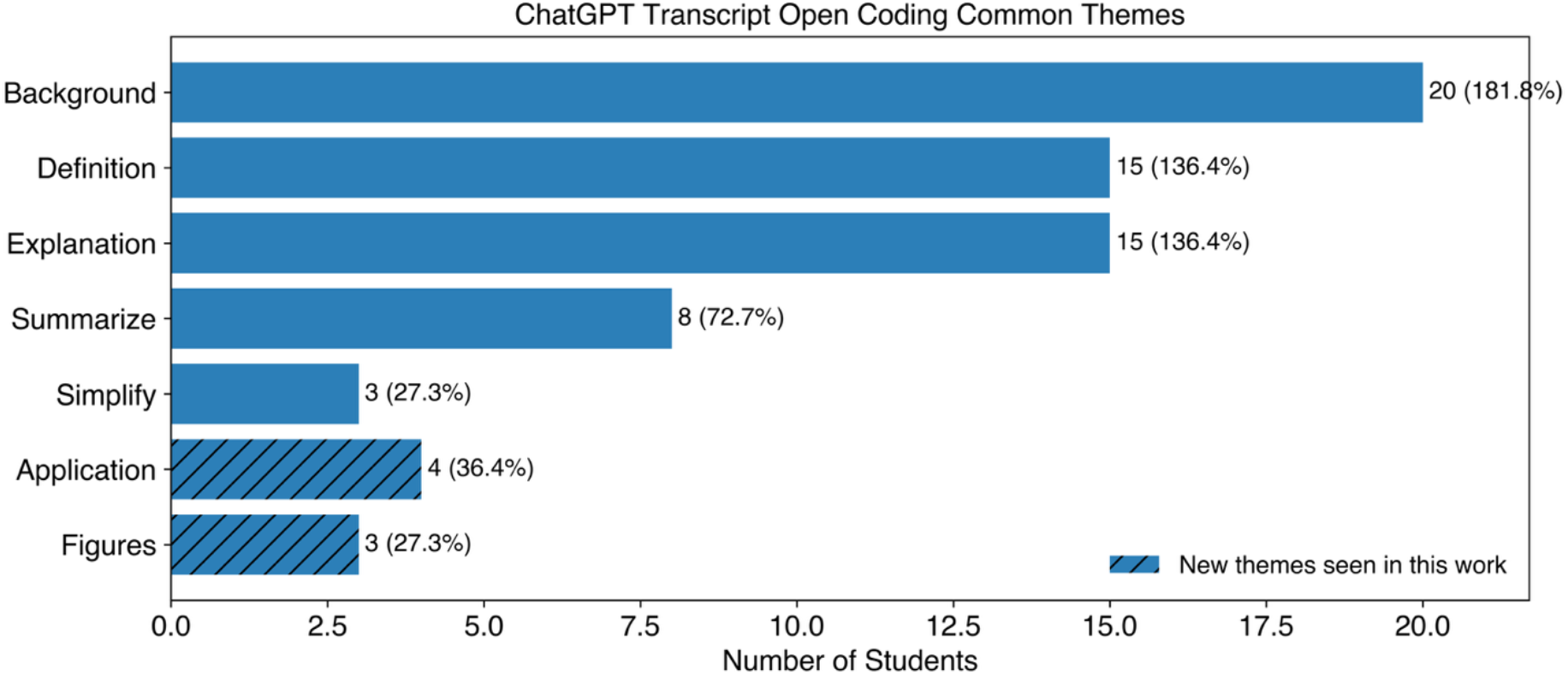
ChatGPT transcript open coding common themes.

“Background” refers to the student giving information about their educational background or giving background information about the journal article being read. All but one student (95%, 20/21 students) gave background information, which they were explicitly asked to do in the reading guide to allow ChatGPT to tailor its response to the student. Students shared their education, research, and/or scientific reading background, and most gave either the title of the journal article being read or the subject. As discussed in Haraldsrud and Odden’s paper,^12^ giving sufficient context leads to more productive interactions with ChatGPT. For example:

> “Context: I am a 3rd year undergraduate biochemistry major with a moderate level of understanding of computational chemistry. I’m going to ask clarifying questions about the article titled “Accurate prediction of protein structures and interactions using a three-track neural network.”

“Definition” and “explanation” were both the next most commonly used type of interaction (71%, 15/21 students), which are considered productive uses of generative AI. Definition interactions involved asking for the definition of an unfamiliar term. For example:

> “What is a three track neural network” and “What is RoseTTAFold?”

Following definition questions, students frequently asked follow-up explanation questions to further clarify their understanding of the unfamiliar term. For example:

> “Could you elaborate and clear up the idea of blind structure prediction tests?”

Students also asked for explanations independently of definition questions to gain deeper insights into concepts presented in the text:

> “The journal includes comparisons of the ‘TM-scores’ for multiple prediction models. Can you give more context to what a TM score is and why it is important?”

The next most common type of interaction was “summarize” in which the student asked for a summary of either the entire text or part of the text (33%, 7/21 students). Of students who requested a summary, 6 of the students gave the citation or PDF file of the paper and asked ChatGPT to summarize the entire article. One student copied an excerpt from the text and requested that ChatGPT provide a summary of the excerpt. For example:

> “[User gives title and authors of journal article] “What are the main points of this article?”
>
> “[User gives title of article] “Unfortunately it is too difficult for me to understand so please break down this article into more manageable chunks, give me the main topic and points, as well as what use this application has to the world at large. Please be extremely detailed and give me definitions for the more difficult words and phrases.”

This use of ChatGPT to provide summaries of the text is one that is of potential concern, as students may be relying on AI-generated summaries rather than reading the literature themselves. This is a non-productive use of the technology, as students may be using the technology to avoid engaging with the actual journal article.

The “application” code was an emergent theme in this data that did not appear in the previous study (19%, 4/21 students). In this code, students asked about how the technology described in the journal article could be used in the future. These were “big picture” questions compared to those posed in interactions coded as “explanation” and went beyond the scope of what was suggested in the reading guide. For example:

“Has RoseTTAFold been used in any major drug discoveries within the 21st century?”

“What applications do you see for humans being able to predict protein structure?”

“What implications might this have for understanding protein functions and disease mechanisms?”

The least commonly used codes were “simplify” and the emergent new theme of “figures” (14%, 3/21 students). The simplify code refers to either asking for simplification of the language from the text of the article or asking ChatGPT to simplify its responses. In this study, one student asked for simplification of text from the article:

“Please simplify the following paragraph: [user copied excerpt from the article].”

The other two students requested that ChatGPT simplify the language used in its responses:

“For future answers keep the *[sic]* brief.” and “I have limited proficiency in understanding scientific journals and articles. So, please try to explain the article in a way I can understand.”

Lastly, the emergent code “figures” refers to the student specifically asking for explanation or description of one or more figure in the text (14%, 3/21 students). Here, students either asked for a description of a specific figure or asked for an outline of all the figures in the article. For example:

“Could you describe/summarize the first figure about comparison of multi track models?” and “Can you further explain the images within the article?”

While most interactions were superficial (requesting definitions and summaries), students did engage in higher-level discussion with ChatGPT in asking for follow-up explanations after definition-type questions, as well as asking about applications of the technology described in the article. However, being able to obtain definitions easily and efficiently through the use of ChatGPT allows students to then engage in higher order thinking and ask follow-up questions. As will be discussed below, most students felt they were able to have richer journal club conversations when they used the ChatGPT-assisted reading guide.

### Student survey/observations from class

Twenty-five (25) students filled out the survey about their experience using the ChatGPT-assisted reading guide. (Note that not every student responded to every question on the survey.) A table of all survey question responses can be found in the Supplemental Information. Overall, most students reported a positive experience with the LLM-assisted reading guide, with 88% of students saying that they would use ChatGPT in the future to read journal articles (**Figure 2)**.

**Figure 2.**
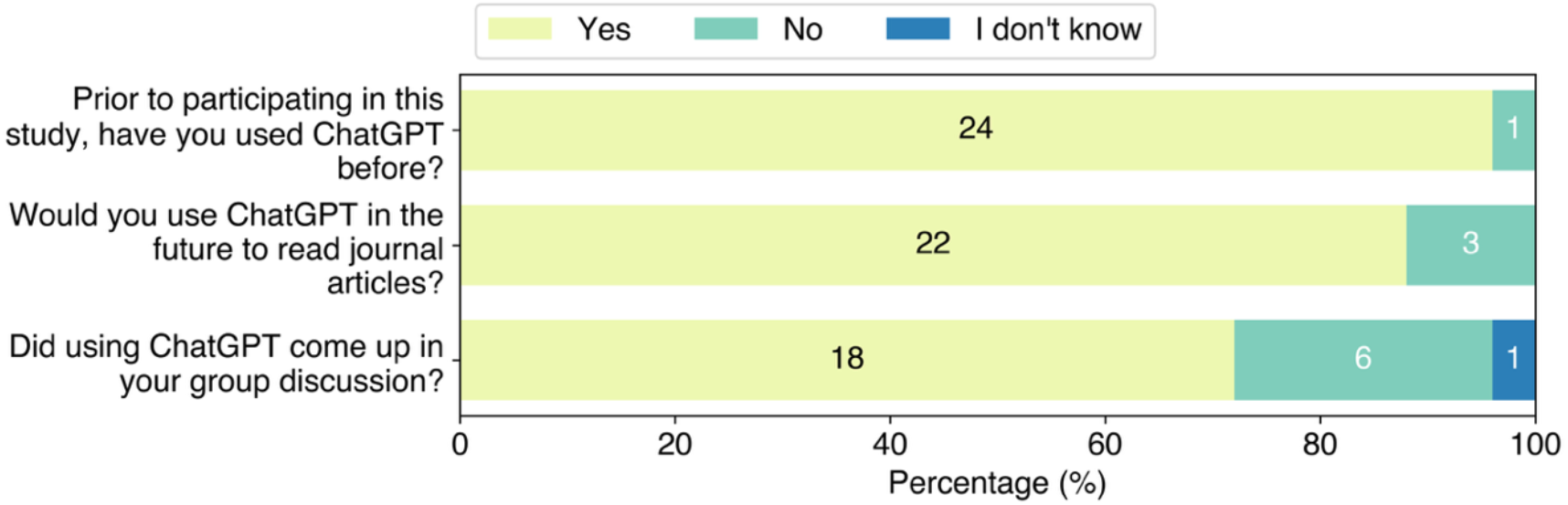
Experience with ChatGPT reported in the student survey.

The overwhelming majority (24 out of 25 students) reported that they had used ChatGPT prior to this assignment (**Figure 2**); commonly cited uses were homework and studying and writing, revising, or outlining written work. Small numbers of students (3 or fewer) reported using LLMs for non-academic uses, coding, or summarizing. Further, students reported a variety of experience in reading scientific literature prior to this study. **Figure 3** show survey data indicating that most students had limited scientific reading experience prior to the study, with most students (21/25) reading two or fewer articles per month. **Figure 4** tabulates reading approaches used prior to the study as reported on the survey, with the most common approach being reading the abstract first. **Figure 5** shows strategies students reported using when encountering unfamiliar terminology prior to this study. The most common reported strategy was using a search engine like Google (95% respondents), followed by using ChatGPT (52%) or skipping the unfamiliar term (52%).

**Figure 3.**
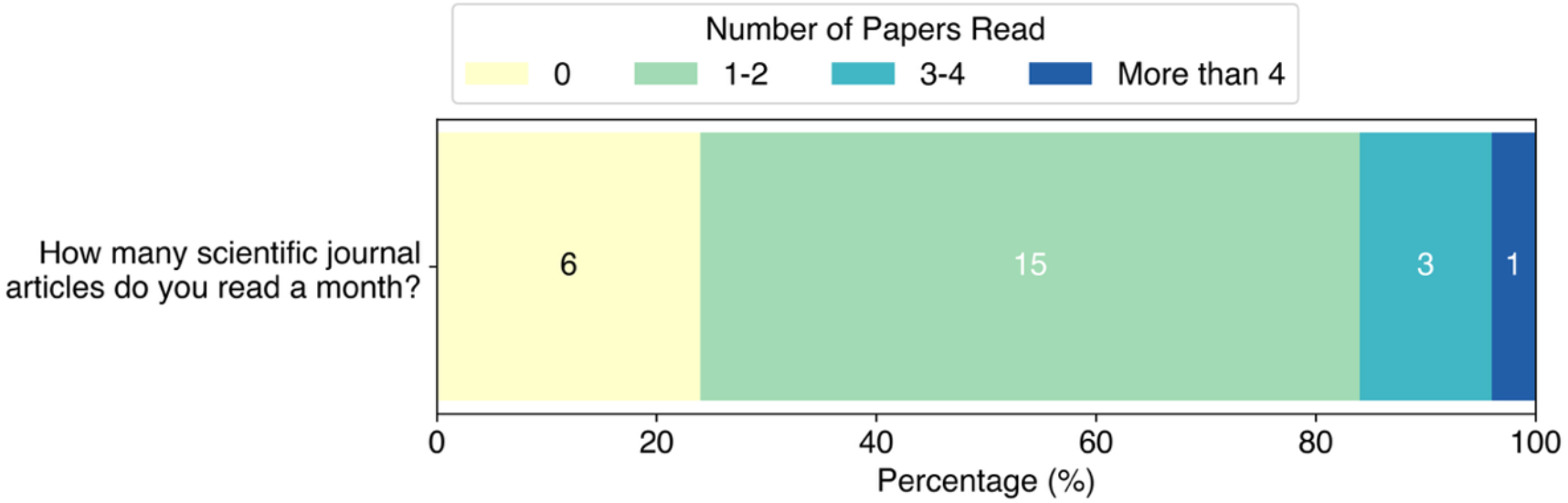
Number of scientific articles read monthly reported in the student survey.

**Figure 4.**
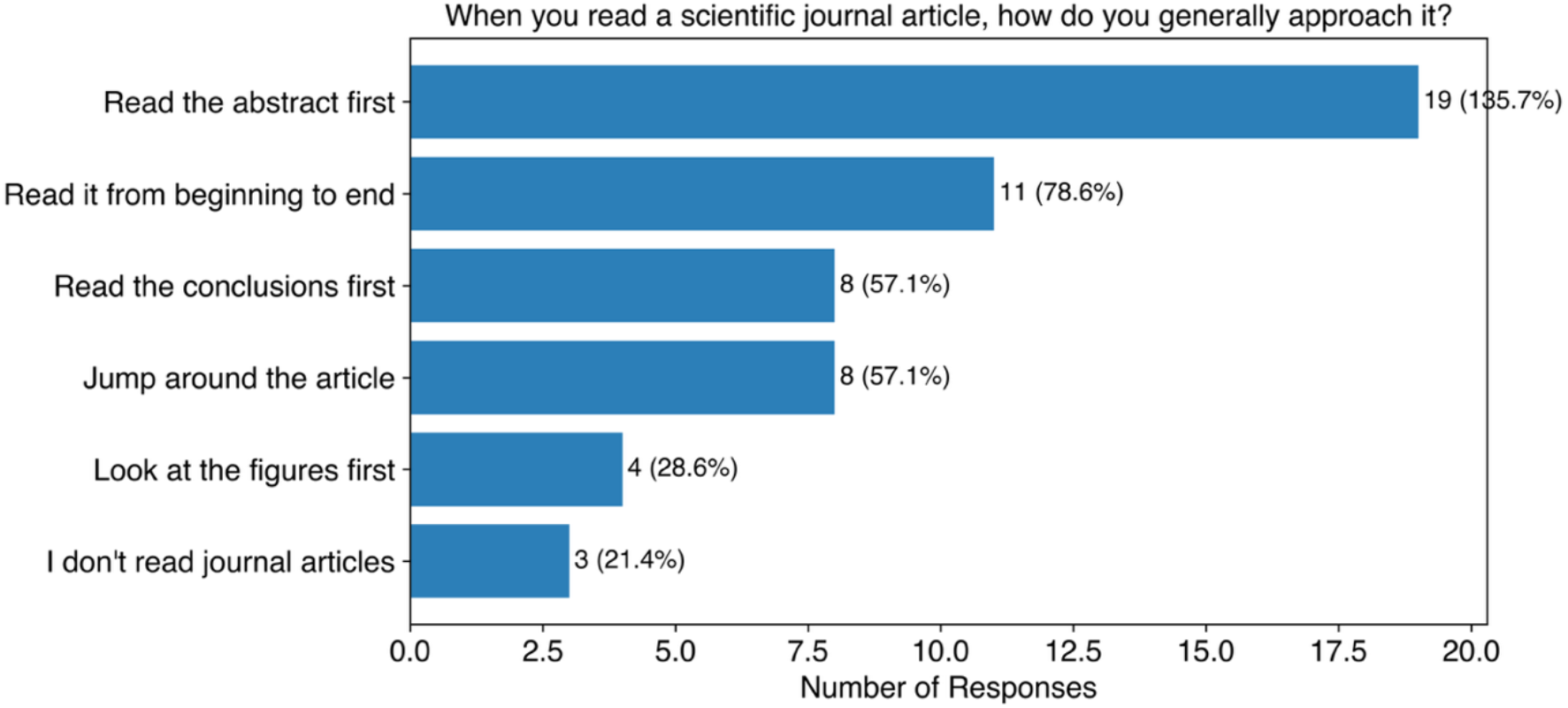
Scientific reading approaches reported in the student survey.

**Figure 5.**
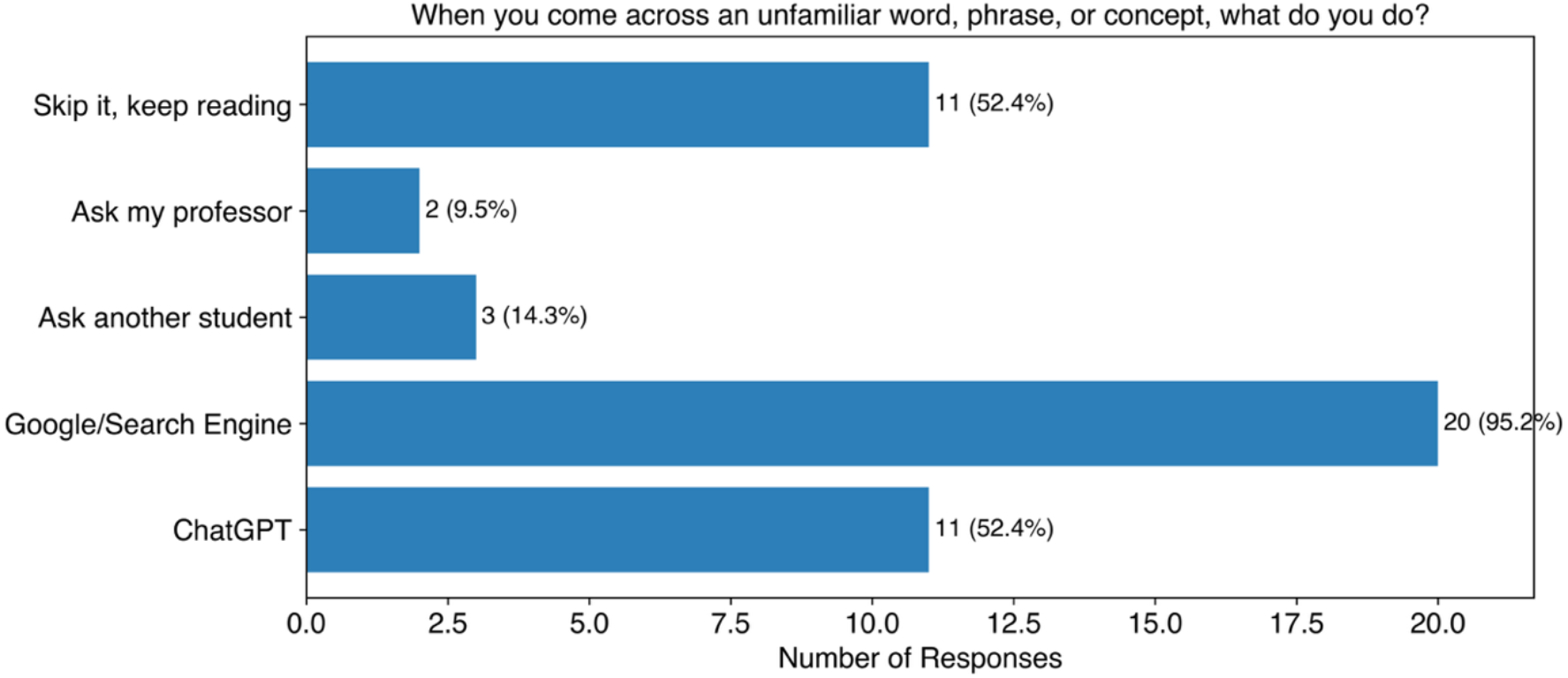
Approaches to unfamiliar terminology in scientific literature reported in the student survey.

Students highlighted several benefits of using ChatGPT, the majority of which were themed around helping them understand unfamiliar concepts. Students highlighted being able to get definitions for unfamiliar language (8 responses), the clarity of explanations provided (5), the ability to ask for summarized information (5), and context about the article (5). Two students specifically mentioned that ChatGPT provided them information that specifically fit their background knowledge, and two students said ChatGPT helped them understand the big picture impact of the study.

There were a few very notable exceptions to the overall positive experiences the students reported. One student in particular, highlighted that they did not feel ChatGPT gave accurate information and should not be used to understand scientific literature because of the likelihood of hallucinations or other incorrect information. (Note that the authors of this study did not in any way attempt to verify or validate any of the information from the ChatGPT transcripts; we were strictly concerned with what types of things the students asked ChatGPT.) In all, five students made comments about their concern about the accuracy of the information ChatGPT provided them. Students also expressed frustration with not getting the response they wanted based on their inquiry (8 responses), complaints that even when they asked for simplified information, the explanations were too hard to understand (4) or that the explanations contained too much information (3). Three students also specifically said that ChatGPT was not useful for learning, with one student saying,

“Chat GPT did not help me learn how to read future articles, nor conditioned me to the same sintax. In a sense, it acts like a crutch that doesnt help with future projects, but is great for the current moment.”

Survey responses about the experience of using ChatGPT in reading the journal article are summarized in **Table 1**.

**Table 1.**
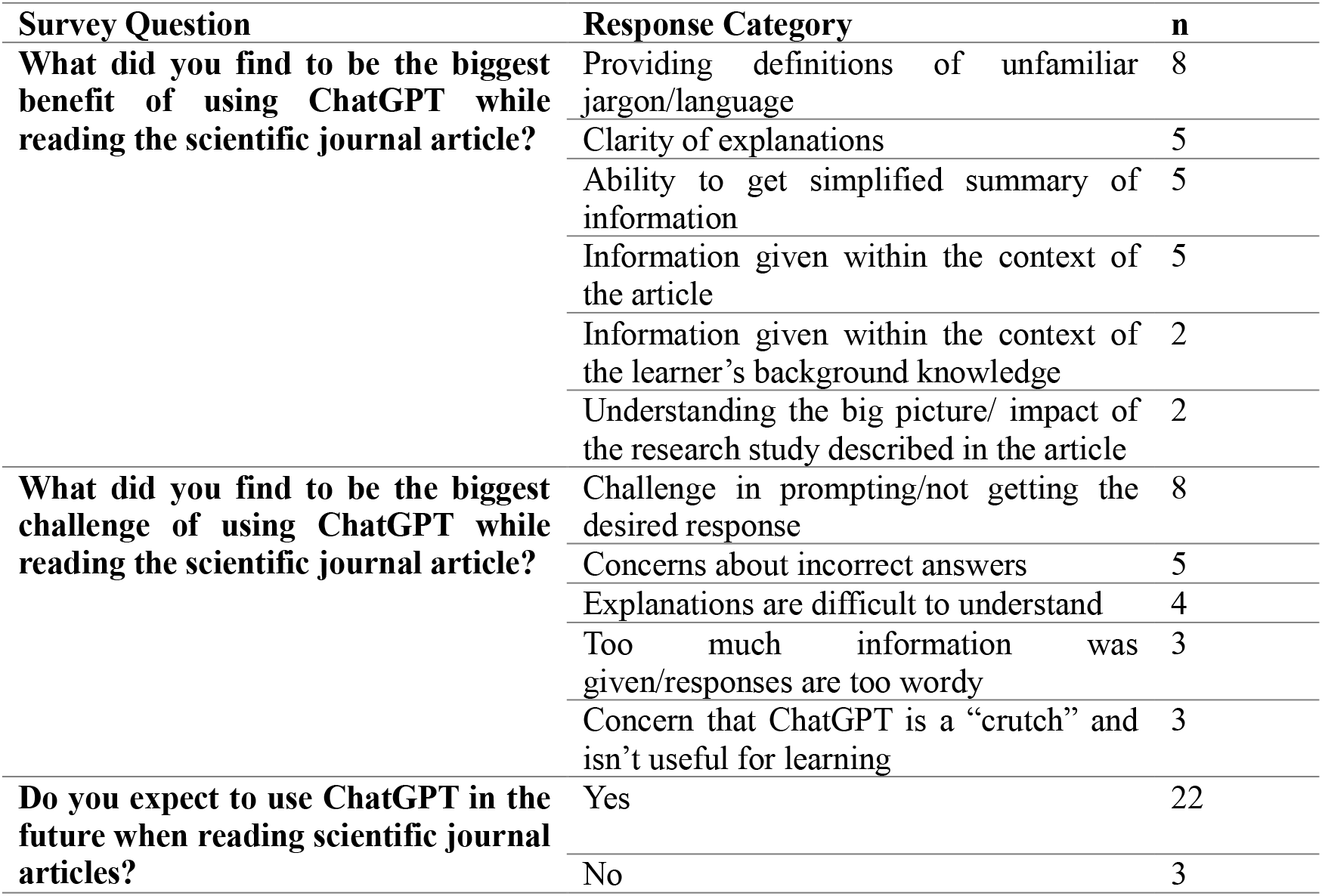
Selected survey responses regarding using the ChatGPT-assisted reading guide to read the scientific journal article.

Anecdotally, the instructor (McDonald) felt that the small group discussions were more lively and more sustained, the week that students had read the paper using the ChatGPT-assisted reading guide. In the first iteration, the instructor had allotted 25 minutes for small group discussion; they observed that after 20 minutes, discussion in most groups was lagging or the group had shifted to discussing something off topic. In the second iteration, the discussion was still very active when time was called after 25 minutes. Participation in the discussions also seemed more balanced, with every student in the group contributing at least some to the discussion. On the survey, students were asked whether their in-class discussion with their peers was better for the paper they had used the ChatGPT-assisted reading guide or the paper where they had not. Thirteen (13) students reported their discussion was better for the paper where they had used ChatGPT; five (5) students said it was better where they had not used ChatGPT, and seven (7) students said there was no difference in the quality of the discussion. The comments also captured these differing viewpoints, with one student saying,

“For the discussion without the use of Chat GPT, I felt like the discussion wasn’t productive since no one really actually understood what was going on in the article and we mostly spent time trying to understand what it meant. In the discussion with the use of Chat GPT, I felt it was a lot more productive since everyone seemed to understand the article and what it was about a lot better, and our discussion was more high-level because if this.”

But another student, in response to the same prompt said,

“I think without the option of AI that people spent more time reading it themselves and thus had a better understanding of the paper.”

Survey responses about the effect of using ChatGPT on their journal club discussions are summarized in **Table 2**.

**Table 2.**
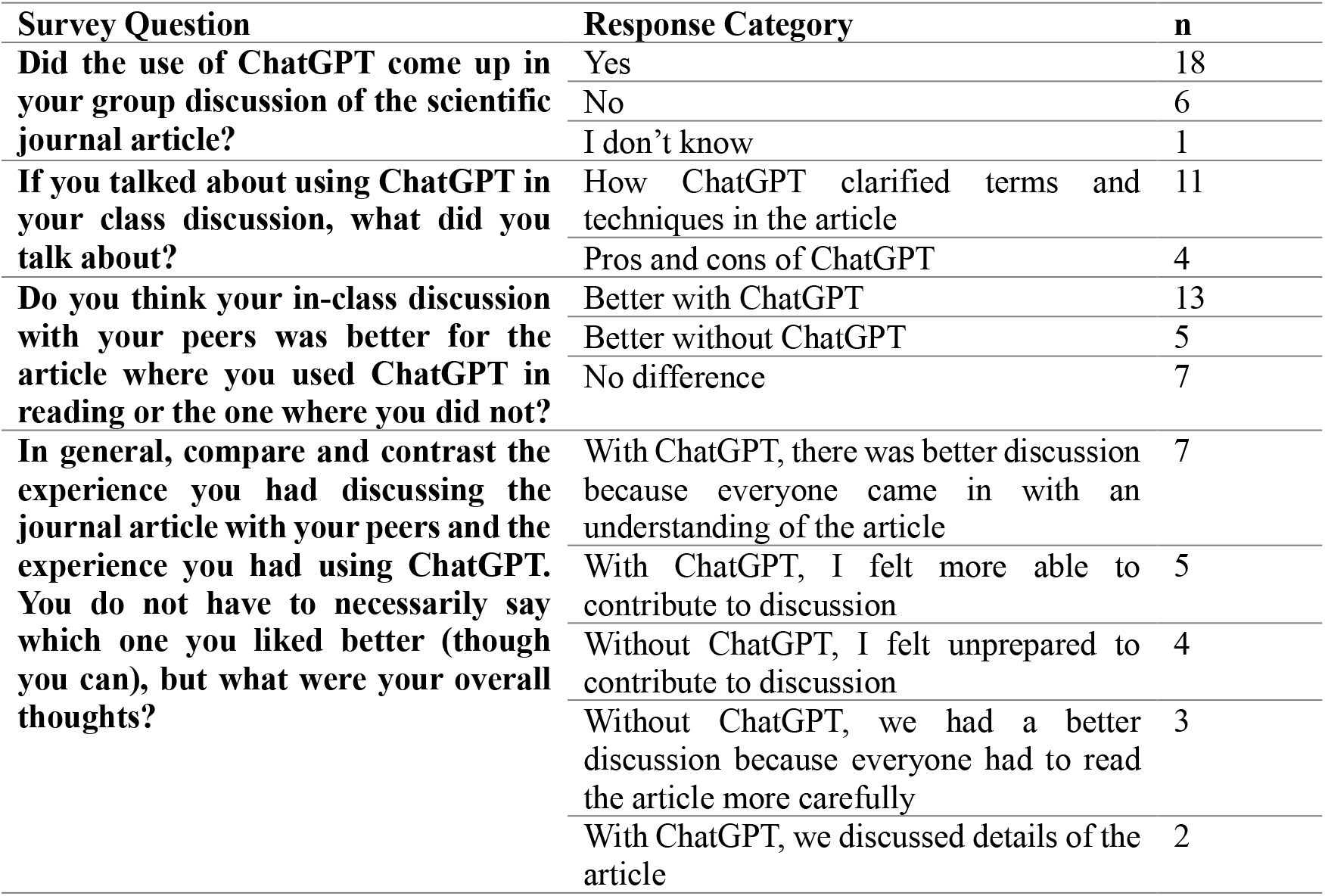
Selected survey responses regarding the effect of the ChatGPT-assisted reading guide on journal club discussions.

### Comparison to initial study

In the initial study of how students used the ChatGPT-assisted reading guide, a small number of undergraduate research students were asked to use the reading guide to read the title, abstract, and introduction to a structural biology journal article.^13^ These sections of the article were chosen for the initial study because this is where students are most likely to first encounter unfamiliar terms and jargon but these sections lacked figures for the reader to interact with. ChatGPT transcripts collected and coded using open iterative coding to find the five interaction themes described above. A pre-survey asked students about their experience with scientific reading. A post-survey asked students about their experience using the ChatGPT-assisted reading guide and experience using ChatGPT.

Many similarities emerge between the initial study and the current study described here. First, most undergraduates have little experience reading scientific journal articles with most reporting reading two or fewer articles per month, and students reported either not being taught reading strategies, or being taught a variety of different strategies. However, both groups of students reported confidence in reading scientific literature that did not match their lack of experience. This highlights the need for improved, sustained reading instruction for undergraduate science students. Both groups of students predominantly relied on search engines such as Google when encountering an unfamiliar term or concept, with about half of students using generative AI such as ChatGPT prior to using the ChatGPT-assisted reading guide. Overall, both groups reported that the reading guide to be effective in helping them read the assigned paper.

However, there were some noticeable differences in how the two groups interacted with ChatGPT and how they interpreted their experience using generative AI. The initial and current study occurred about a year and a half apart from each other, and during that time generative AI and ChatGPT became more ubiquitous in both academia and everyday life and students gained more experience with using ChatGPT during that time. This led to some differences in how the students in the initial study and the current study interacted with ChatGPT. Most noticeably, students in the current study asked “summarize” type questions more often than students in the initial study. Further, the “summarize” interactions in the current study frequently involved students asking ChatGPT to summarize the entire paper, possibly in lieu of reading the paper. This is not an effective or productive use of the technology, because students can use it to avoid reading a paper rather than using it to clarify questions while actively engaging with the paper. Classroom instruction on scientific reading using the ChatGPT-assisted reading guide should emphasize the importance of active reading and using ChatGPT for clarification rather than using ChatGPT to completely avoid struggling with reading scientific literature.

In the initial study, student feedback about using ChatGPT to read the article was overwhelmingly positive, with no students expressing concern about the veracity of the information or other potential negative impacts of using generative AI. However, with more experience, students in the current study expressed more skepticism of the use of generative AI in the survey conducted after the journal club activities. Multiple students expressed concerns about the trustworthiness of the information provided by ChatGPT, most likely due to other experiences they had with ChatGPT. Further, some students shared concerns about the environmental impact of generative AI. Lastly, multiple students shared concerns about overreliance on AI in their education and expressed concern that the use of AI can affect their own learning. While most students felt more confident in their reading after using ChatGPT and felt that their journal club discussions were more productive, some students also expressed concerns about overreliance on the technology. This shows that students are developing a more nuanced view of generative AI with experience using it in various applications.

## Conclusions/recommendations

Here we discuss the use of a ChatGPT-assisted reading guide to support students in a journal club assignment in a chemistry seminar course. Students engaged in two in-class journal club activities; they did not use generative AI for the first assignment, and they used the ChatGPT-assisted reading guide in the second assignment. Students’ ChatGPT transcripts were analyzed for common types of interactions, based on themes observed in an initial study of using the ChatGPT-assisted reading guide. We found that students primarily used ChatGPT to obtain definitions and explanations of unfamiliar terminology and concepts. Unlike in the initial study, many students also used ChatGPT to summarize the article, possibly in lieu of reading the article in detail. Further, themes of questions about applications of the technology described in the article and questions about article figures emerged in this study. Students completed a survey about their experience after the assignments. The survey responses indicated that the students are thinking critically about the implications of generative AI use in their education. Even among students who said they would use ChatGPT in the future, students highlighted several important issues including the validity of the information provided by ChatGPT and how using generative AI affected their own thinking, reasoning, and creativity. If faculty wish to incorporate the use of generative AI in their curriculum, we recommend they address these issues directly with their students, discussing the benefits and shortfalls of generative AI,^15^ as well as ethical use of generative AI,^16^ in the context of how it is being used in their class.

## Supporting information

Supplemental Information: ChatGPT-Assisted Reading Guide and Survey Data

