## Supplemental Information: ChatGPT-Assisted Reading Guide and Survey Data for "Benefits and Challenges of Integrating a Generative AI Assisted Reading Guide in an Undergraduate Journal Club Assignment"

1. The ChatGPT-Assisted Reading Guide

### Guidelines for Reading Scientific Papers

**Objective:** The primary aim of this method is to facilitate a smooth reading experience, minimizing disruptions to your flow. The preview stage primes you for the paper's content, while the reading stage aims to maintain a steady pace. The use of ChatGPT serves as a real-time resource for overcoming stumbling blocks in comprehension.

##### Previewing the Paper

1. **Understand the Title**: Determine the main focus of the paper based on its title.
2. **Analyze the Abstract**: Read the abstract, and if available, the accompanying figure, to identify the paper's major findings.
3. **Abstract Structure**: While reading the abstract, consider the typical structure: Importance, Problem, Approach, and Insight.
4. **Review Headings and Figures**: Skim through the paper's headings and figures to anticipate the questions the paper aims to answer.

##### Reading the Paper

1. **Highlight Key Text**: Use an erasable method (e.g., pencil underlining) to highlight text that seems important or intriguing. You can refine this later.
2. **Margin Notes**: Whenever you have a question, insight, or idea, jot it down in the margins. These notes are for your personal use.
3. **Extended Notes**: If you need more space for notes or sketches, use a separate notebook or sheet of paper. The goal is to capture all your thoughts for later reference.

##### Utilizing ChatGPT for Clarifications

4. **When you encounter an unfamiliar concept, term, or methodology that feels like a stumbling block, use ChatGPT to help clarify the concept**:

a. **Start a New Conversation**: Initiate a new conversation with ChatGPT. Use this same thread for all questions related to this paper.
b. **Introduce Yourself**: Briefly describe your academic background, lab experience, and comfort level with the material and reading scientific papers.
c. **Inform ChatGPT About the Paper**: Provide the paper's title, field, and any other relevant information to help ChatGPT understand the context.
d. **Ask Your Questions**: Pose your question to ChatGPT. If the initial response is unclear, continue asking follow-up questions until you get the information you're seeking.
e. **Seek Further Clarification**: If something remains unclear, ask for additional context or a different type of explanation.
f. **Explore Further**: If a point seems particularly relevant or interesting, feel free to continue the conversation with ChatGPT as long as needed to fully understand the paper.

5. **Post-Reading Summary**: After completing the paper, review your notes and write a brief summary that you can refer to later. This can be as short as one sentence.

2. Full survey data

| Survey Question | Response Category | n |
|---|---|---|
| **If you have been taught strategies for reading scientific journal articles, what strategies were you taught?** | None | 7 |
| | Read the abstract, conclusions, figures, then rest of paper | 7 |
| | Skim the paper and then read it in depth | 6 |
| | Use Google or AI to cross-reference | 2 |
| | Read the figures and ID key words | 2 |
| **When you read a scientific journal article and come across an unfamiliar word, phrase, or concept, what do you do?** | Look it up with a search engine like Google | 20 |
| | Skip it and keep reading | 11 |
| | Look it up with AI | 11 |
| | Ask another student | 2 |
| | Ask a professor | 2 |
| **What do you think is the most challenging part of reading scientific journal articles?** | Unfamiliar, specialized knowledge | 14 |
| | Language/jargon/vocab | 12 |
| | Dense writing | 3 |
| | Parsing out the important information | 3 |
| **Do you feel confident reading scientific journal articles?** | 1 (not at all confident) | 1 |
| | 2 | 7 |
| | 3 | 9 |
| | 4 | 7 |
| | 5 (extremely confident) | 1 |
| **How many scientific journal articles do you read per month?** | 0 | 6 |
| | 1 – 2 | 15 |
| | 3 – 4 | 3 |
| | More than 4 | 1 |
| | Yes | 24 |

| | | |
|---|---|---|
| **Prior to this study, have you used ChatGPT before?** | No | 1 |
| **If you have used ChatGPT before this study, what did you use it for?** | Homework and studying | 14 |
| | Writing, revising, outlining, and/or planning essays | 6 |
| | Summarizing information | 3 |
| | Non-academic uses | 3 |
| | Reviewing work for errors | 2 |
| | Using ChatGPT as a general search engine | 2 |
| | Coding | 2 |
| **What did you find to be the biggest benefit of using ChatGPT while reading the scientific journal article?** | Providing definitions of unfamiliar jargon/language | 8 |
| | Clarity of explanations | 5 |
| | Ability to get simplified summary of information | 5 |
| | Information given within the context of the article | 5 |
| | Information given within the context of the learner's background knowledge | 2 |
| | Understanding the big picture/ impact of the research study described in the article | 2 |
| **What did you find to be the biggest challenge of using ChatGPT while reading the scientific journal article?** | Challenge in prompting/not getting the desired response | 8 |
| | Concerns about incorrect answers | 5 |
| | Explanations are difficult to understand | 4 |
| | Too much information was given/responses are too wordy | 3 |
| | Concern that ChatGPT is a "crutch" and isn't useful for learning | 3 |
| **Do you expect to use ChatGPT in the future when reading scientific journal articles?** | Yes | 22 |
| | No | 3 |
| **Did the use of ChatGPT come up in your group discussion of the scientific journal article?** | Yes | 18 |
| | No | 6 |
| | I don't know | 1 |
| **If you talked about using ChatGPT in your class discussion, what did you talk about?** | How ChatGPT clarified terms and techniques in the article | 11 |
| | Pros and cons of ChatGPT | 4 |
| **Do you think your in-class discussion with your peers was better for the article where you used ChatGPT in** | Better with ChatGPT | 13 |
| | Better without ChatGPT | 5 |
| | No difference | 7 |

| | | |
|---|---|---|
| **reading or the one where you did not?** | | |
| **In general, compare and contrast the experience you had discussing the journal article with your peers and the experience you had using ChatGPT. You do not have to necessarily say which one you liked better (though you can), but what were your overall thoughts?** | With ChatGPT, there was better discussion because everyone came in with an understanding of the article | 7 |
| | With ChatGPT, I felt more able to contribute to discussion | 5 |
| | Without ChatGPT, I felt unprepared to contribute to discussion | 4 |
| | Without ChatGPT, we had a better discussion because everyone had to read the article more carefully | 3 |
| | With ChatGPT, we discussed details of the article | 2 |
